# Iterative point set registration for aligning scRNA-seq data

**DOI:** 10.1101/2020.05.13.093948

**Authors:** Amir Alavi, Ziv Bar-Joseph

## Abstract

Several studies profile similar single cell RNA-Seq (scRNA-Seq) data using different technologies and platforms. A number of alignment methods have been developed to enable the integration and comparison of scRNA-Seq data from such studies. While each performs well on some of the datasets, to date no method was able to both perform the alignment using the original expression space and generalize to new data. To enable such analysis we developed Single Cell Iterative Point set Registration (SCIPR) which extends methods that were successfully applied to align image data to scRNA-Seq. We discuss the required changes needed, the resulting optimization function, and algorithms for learning a transformation function for aligning data. We tested SCIPR on several scRNA-Seq datasets. As we show it successfully aligns data from several different cell types, improving upon prior methods proposed for this task. In addition, we show the parameters learned by SCIPR can be used to align data not used in the training and to identify key cell type-specific genes.

**Author Summary:** Integrating single cell expression data (scRNA-Seq) across labs, platforms, and technologies is a major challenge. Current methods for addressing this problem attempt to align cells in one study to match cells in another. While successful, current methods are unable to learn a general alignment in *gene space* that can be used to process new or additional data not used in the learning. Here we show that the scRNA-Seq alignment problem resembles a well known problem in the field of computer vision and robotics: point-cloud registration. We next extend traditional iterative rigid-object alignment methods for scRNA-seq while satisfying a set of unique constraints that distinguishes our solution from past methods. Analysis of transcriptomics data demonstrates that our method can accurately align scRNA-seq data, can generalize to unseen datasets, and can provide useful insights about genes active in the cells being studied.

## 1 Introduction

While only recently introduced, single-cell RNA-sequencing (scRNA-seq) has quickly developed into an indispensable tool for transcriptomics research. Driven by the development of droplet microfluidics-based methods [1, 2, 3, 4], current experiments are able to simultaneously profile expression of genes in tens of thousands of single cells. Studies ranging from cell type and state identification [5, 6] to tracking early development [7, 8] to unveiling the spatial organization of cells [9, 10] are all utilizing scRNA-Seq data, providing new insights about the activity of genes within and between cells.

While the size and number of individual scRNA-seq datasets is large and constantly growing, the question of how to integrate scRNA-Seq data from multiple experiments or platforms has become increasingly relevant. Different labs are seeking to analyze related tissues in an organ system, such as mapping out the cell types in the human pancreas [11] or building an adult mouse brain cell atlas [12]. On on even larger scale, consortia such as the Human Cell Atlas [13, 14] or the HUBMaP [15] are organizing researchers globally with the goal of mapping cells in the entire human body.

Combining datasets, even for the same tissue, across platforms or labs is a challenging problem. This process is often referred to “dataset alignment”, “dataset harmonization”, or “batch correction,” and is an active area of research. A number of methods have been recently suggested to address this problem. Many of these rely on nearest neighbors computations. For example, Mutual Nearest Neighbors (MNN) integrates two datasets by first identifying cells in the two datasets that are mutual nearest neighbors (in each other’s set of *k* nearest neighbors) [16]. It then computes vector differences between these pairs and uses weighted averages of these vector differences to shift one batch onto the other. Another method, Seurat [17], extends this idea by first computing MNNs in a reduced dimension space, via canonical correlation analysis (CCA) which identifies common sources of variation between the two datasets, and then proceeding to correct the batch effects in a similar fashion as MNN. Other methods such as scVI [18] and ScAlign [19] use a neural network embedding to align the two datasets. These methods seek to encode the scRNA-seq datasets using a common reduced dimensional space in which the batch effects are reduced. While the above methods are unsupervised, there are also a few supervised methods proposed for this task. These method require as input the correct cell type labels for cells in the training data and use that to learn a function to assign cell types for the test data. An example of such method is Single Cell Domain Generalization Network (scDGN) which uses a supervised neural network trained with adversarial objectives to improve cell type classification [20]. Another example is Moana, which uses hierarchical cell type classifiers robust to batch effects to project labels from one dataset onto another [21].

Each of the methods mentioned above offers different features and so might be appropriate for different settings. For example, some methods align the data in the given gene space and thus maintain gene semantics while others, namely the neural network-based methods, do the alignment in a new embedded space (i.e. a reduced dimensional space). On the other hand, the neural network methods typically are learning an alignment function which enables the alignment to be applied to new data (generalization). A comparison of the features of each of the methods is summarized in Table 1.

**Table 1:**
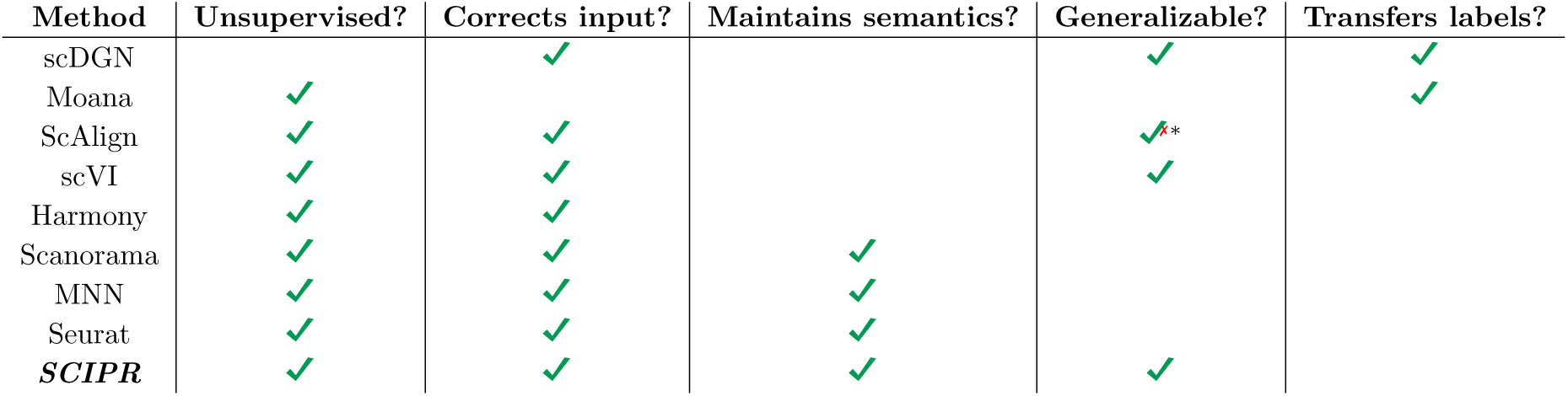
Comparison of features and properties of various scRNA-seq alignment methods. The “Corrects input?” column refers to whether the method actually aligns (transforms) the input data batches in order to integrate them. The “Maintains semantics?” column refers to whether the output of the method retains the gene semantics given as input. The “Generalizable?” column refers to whether the method learns a model which can be applied to new data. The “Transfers labels?” column refers to whether the method also explicitly aims to apply the cell type labels of one data batch onto another, unlabeled batch. * ScAlign is theoretically able to be applied on new data, as it learns a neural network embedding model, but the ability to save and load the function in different sessions to apply it on new data was not available in software at the time of testing.

As the table shows, non of the current methods enables both, maintenance of semantics (required for analyzing genes following the alignment) and generalization (required for keeping the alignment consistent when new data arrives). Here we propose a new method, Single Cell Iterative Point set Registration (SCIPR), which achieves both using an unsupervised framework. Our method extends a well known method in image analysis termed iterative closest points (ICP), which is used for the problem of point-set or point-cloud registration [22]. In ICP, two datasets are represented as sets of points in a common coordinate system, and the method proceeds by pairing together points between the two sets and learning a transformation to move one set closer to (the corresponding points) in the other [23].

We tested SCIPR on three benchmark datasets and compared its performance to several prior methods suggested for the alignment task. As we show, single cell iterative point set registration outperforms prior methods for most of the tasks and is able to generalize to both unseen data in the target and in the source batch by learning a general function which can be applied to new data. Finally, since it retains the original (gene space) representation, the coefficients learned by single cell iterative point set registration can be used to identify key genes related to the cell types being analyzed.

## 2 Results

### 2.1 Method and benchmarking overview

We developed SCIPR which aligns two batches of scRNA-seq data (termed source and target) using methods motivated by point set registration algorithms. SCIPR first identifies corresponding pairs of cells between source and target batches (Figure 1 panel 1). Rather than using the closest cell (as defined by euclidean distance) in the target to match a source cell, SCIPR uses either of two matching algorithms to account for the heterogeneity and noise in scRNA-seq data: Mutual Nearest Neighbors (MNN) matching [16], and a novel greedy matching algorithm (Algorithm S1, Methods). Once a pairing of cells is established (Figure 1 panel 2), a transformation function is learned to transform source cells so that they are closer to their matched target cell (Figure 1 panel 3). To allow for accurate alignment of high-dimensional scRNA-seq data, we replace the rigid transformation commonly used for point cloud registration with affine transformations. After fitting the transformation function (Methods), we apply it to the source cells (Figure 1 panel 4), and iteratively repeat the process until convergence. The final alignment function we learn is a composition of the transformation functions learned at each iteration (Methods) (Figure 1 panels 5,6).

**Figure 1:**
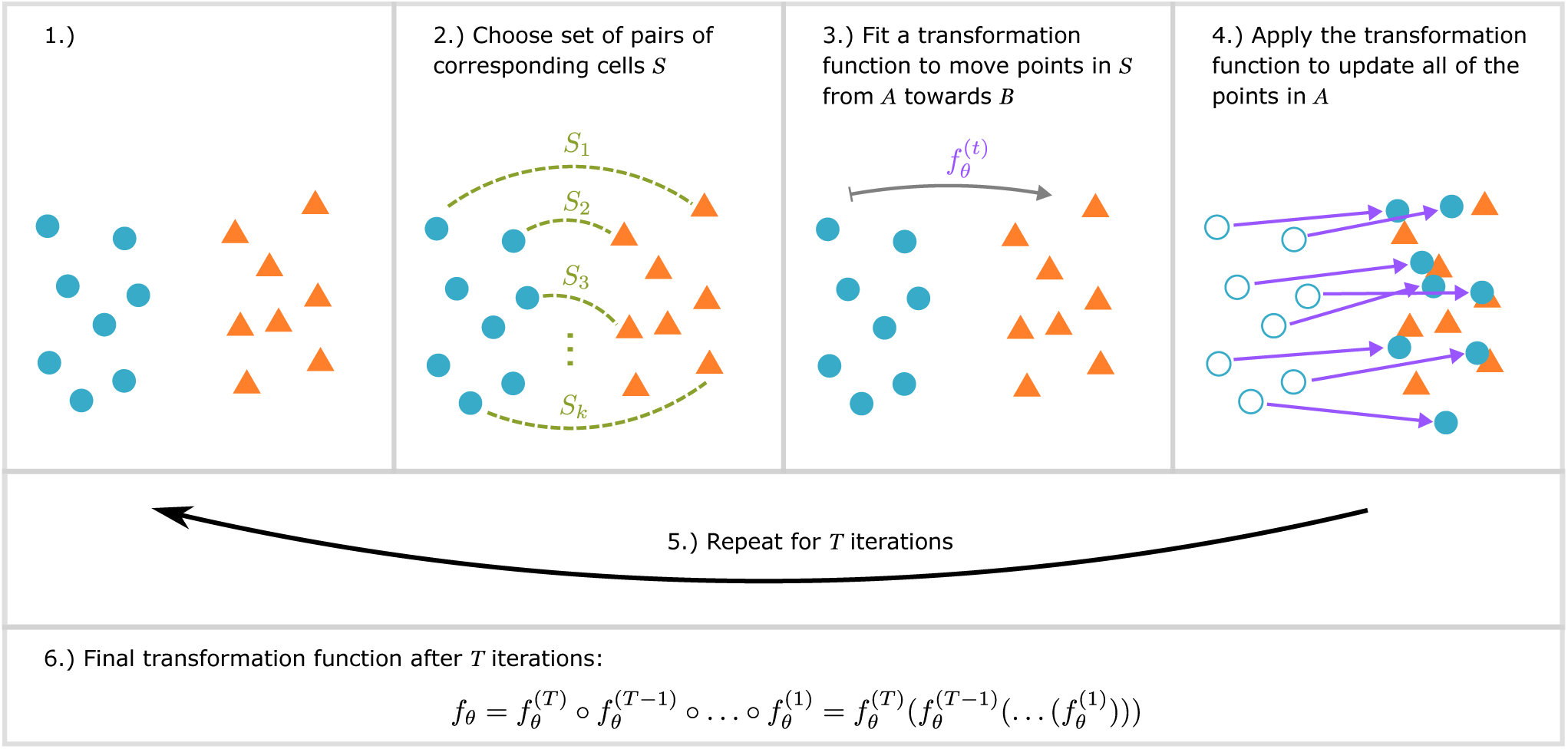
Summary of steps in iterative point set registration for scRNA-seq data. Each cell in an scRNA-seq dataset can be viewed as a point in high dimensional space. 1) We start with two unaligned batches (sources, blue and targets, orange). 2) A matching algorithm (e.g. picking the closest corresponding point, or using mutual nearest neighbors) is used to pair source cells from *A* with a corresponding target cell in *B*. The number of source and / or target cells matched can vary for different matching strategies. 3) Based on the selected pairs, a global transformation function is learned so that source cells in *A* become closer to their paired cell in *B*. 4) The learned transformation is next applied to all points in *A*. 5) This process (steps 2-4) is repeated, iteratively aligning set *A* onto *B* until the mean distance between the assigned pairs of cells no longer improves. 6) The final global transformation function is the composition of the functions learned in each iteration at step 3.

We used three datasets to test and compare two versions of SCIPR to prior alignment methods (Methods). These comparisons were performed by testing the methods on several “alignment tasks”. An alignment task is defined by:

- A dataset (e.g. Pancreas)
- A source batch *A* within that dataset, which you would like to transform (e.g. indrop1)
- A target batch *B* within that dataset, which you would like to transform *A* onto (e.g. indrop 3)

For example, an alignment task can be summarized with the notation: *Pancreas: indrop1* → *indrop3*. In the comparisons we performed we fix the target within a dataset to be the largest batch in that dataset. We scored the performance of the methods using local inverse Simpson’s Index (LISI) in which higher integration LISI (iLISI) is better and lower cell-type LISI (cLISI) is better [24] (Methods).

### 2.2 An affine global transformation function yields well-mixed alignments

We first evaluated the ability of SCIPR and other methods to integrate pairs of batches from three different datasets. Results for 8 alignment tasks in three datasets are presented in Figure 2. As the figure shows, for 7 of the 8 alignment tasks the two version of SCIPR ranked at the top. SCIPR-mnn was the overall top performer ranking first on 4 tasks and 2nd on 2 whereas SCIPR-gdy ranked first on 3 tasks and 2nd on 1. The only other method that performed well is ScAlign which ranked first on 1 task and 2nd on 4. For example, for the CellBench alignment tasks (first row of Figure 2), we see that SCIPR-mnn, which uses the MNN matching for the cell pair assignment stage, has consistent better performance, and achieves high batch mixing (1.70 and 1.76 median iLISI scores on *CELseq2*→*10x* and *Dropseq*→*10x* respectively) with very little cell type mixing (1.00 median iLISI score on both *CELseq2*→*10x* and *Dropseq*→*10x*). When looking at the same dataset, on the *CELseq2*→*10x* task the other methods such as ScAlign (iLISI: 1.00, cLISI: 1.00) or SeuratV3 (iLISI: 1.51, cLISI: 1.00) are also able to avoid cell type mixing, but are not able to mix the batches as much as SCIPR (Figure 2). Full alignment quantitative scores for these tasks and all others in the paper are listed in Table S5. These quantitative metrics are also corroborated by a qualitative assessment (Figure 4). There we can see that both SCIPR-gdy and SCIPR-mnn (top two rows) mix the batches well (1st and 3rd columns) compared to methods like MNN and ScAlign while successfully keeping cell types separate (2nd and 4th columns).

**Figure 2:**
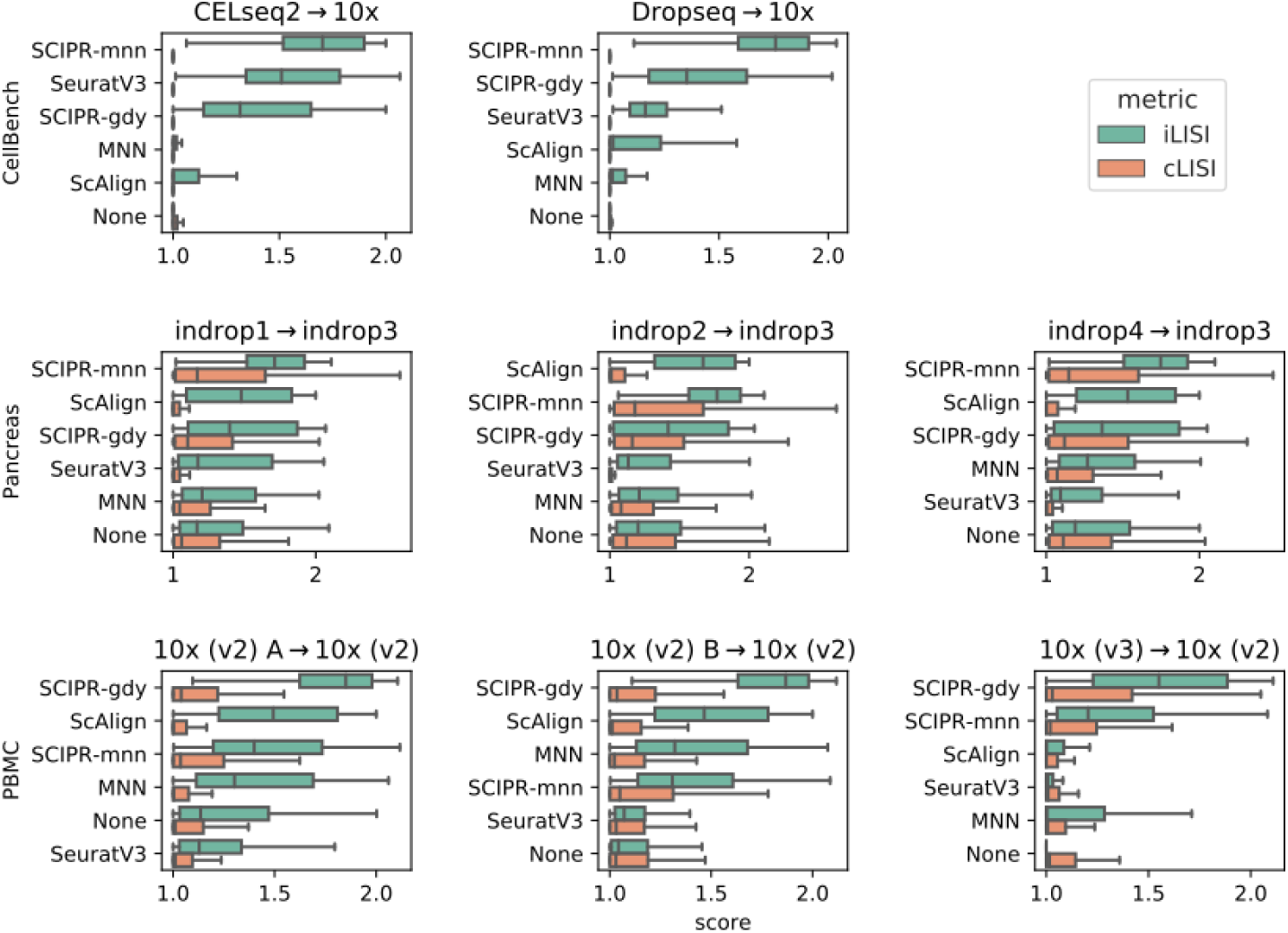
Quantitative scoring of alignment methods on benchmark datasets. Each row of subplots are tasks from the same dataset, where each column uses a different source batch (all are aligned to the same largest reference batch). The scores are iLISI (green, batch integration score), and cLISI (orange, cell type mixing score). In each subplot the methods are ordered from top to bottom in order of largest difference (median iLISI - median cLISI) of scores. The center of each box is the median, and whiskers represent 1.5 times the IQR past the low and high quartiles.

**Figure 3:**
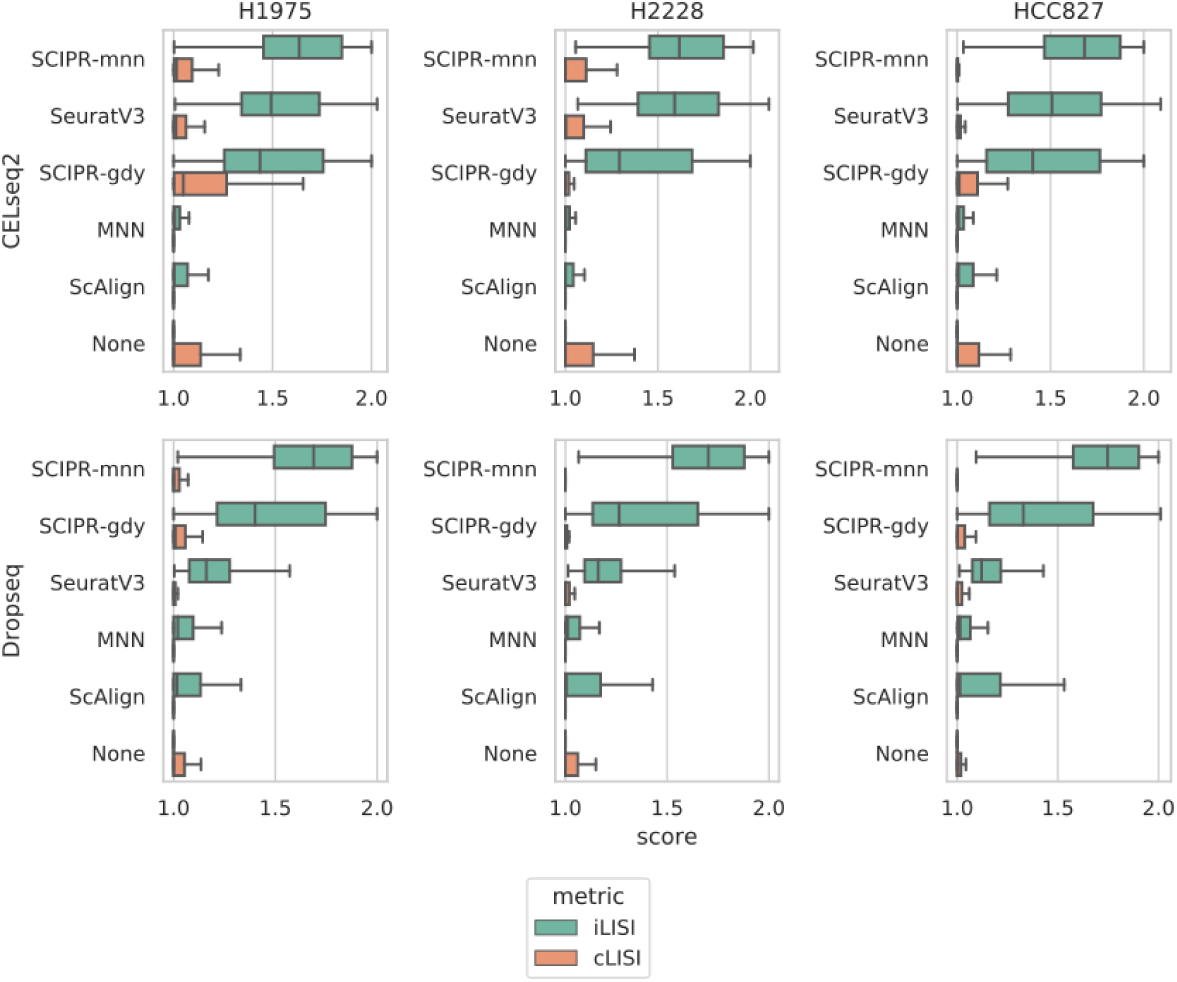
Quantitative scoring of alignment methods on the CellBench dataset with a cell type held out from the target set. Each row of subplots are alignment tasks with the same source batch, where each column uses a different cell type as a hold-out from the target set (the 10x batch). Order and box plot computation are similar to Figure 2.

**Figure 4:**
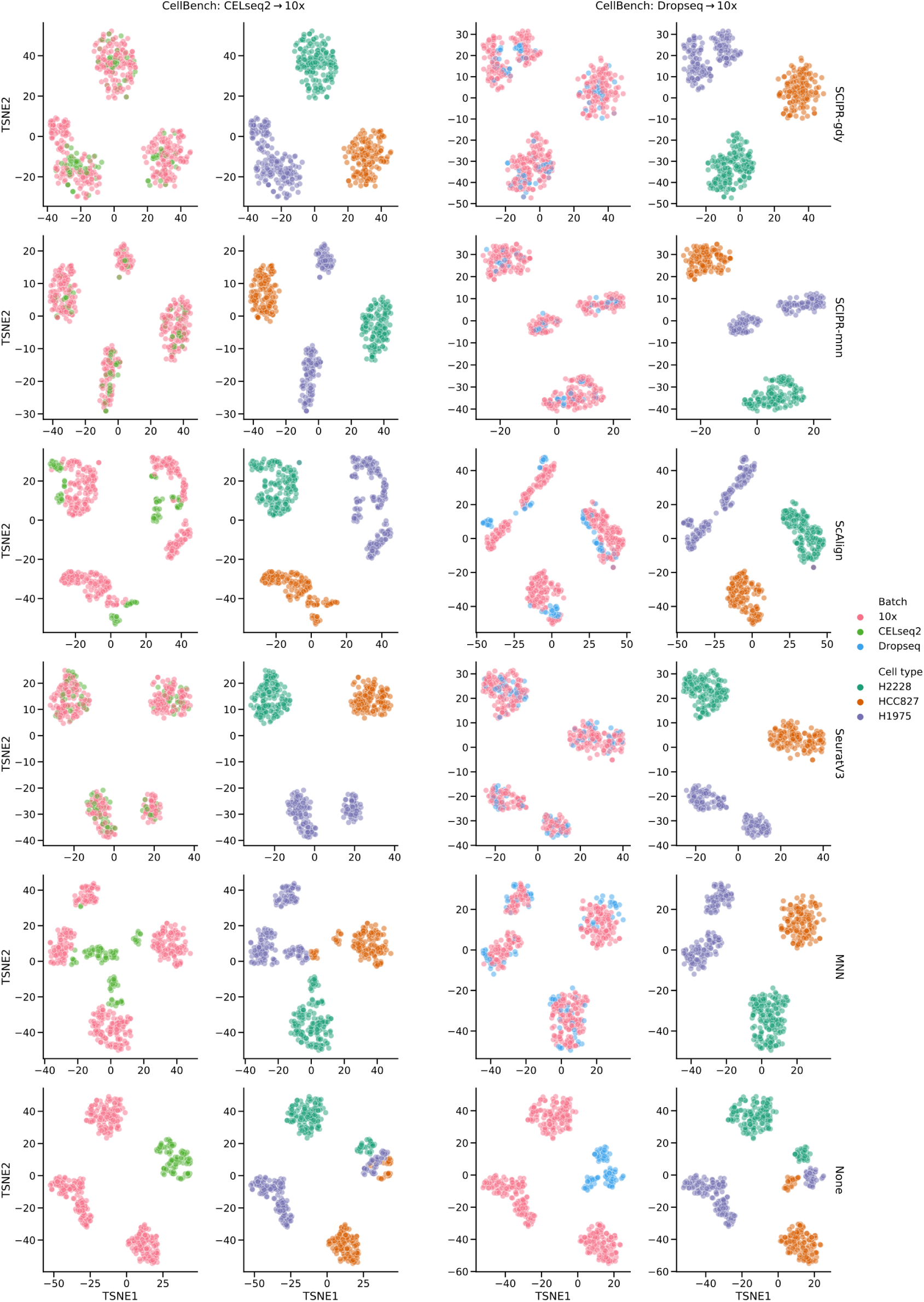
Embedding (t-SNE) visualization from alignment tasks on the CellBench dataset using various alignment methods. Each row is a different alignment method (the bottom row is with no alignment). The columns are in two groups based on alignment task: the left two columns pertain to aligning the CELseq2 batch onto the 10x batch, the right two columns are for aligning the Dropseq batch onto the 10x batch. The first and third columns are colored by batch, and the second and fourth columns are colored by cell type.

### 2.3 SCIPR robustly mixes batches with non-overlapping cell types

The comparisons presented above involved sources and target batches with the same set of cells. However, in practice it is often unknown if both source and target indeed contain the same cell types. To test the robustness of SCIPR and other methods for such realistic scenarios we hold-out a complete cell type from the target set *B* in each of the alignment tasks from section 2.2. As the figures show, for these alignment tasks SCIPR is able to mix batches well, while keeping the median cell type mixing (iLISI) score low, though with a longer tail (Figures 3, S4, S5). For example, for the *CellBench: CELseq2*→*10x (H1975 cell type held-out from target)* task, SCIPR-mnn had median iLISI and cLISI scores of 1.63 and 1.02 respectively while the second best method, SeuratV3, had iLIS and cLISI scores of 1.49 and 1.01 respectively (Figure 3). On the other hand, for the task *Pancreas: indrop1*→*indrop3 (acinar cell type held-out from target)*, SCIPR-mnn achieves a higher median batch mixing score of iLISI=1.69 compared to ScAlign’s score of 1.57, but also mixes the cell types slightly more (SCIPR-mnn median cLISI score: 1.28, ScAlign median cLISI score:1.00) (Figure S4).

### 2.4 SCIPR generalizes to unseen data

One of the advantages of SCIPR compared to most previous methods is the fact that it learns a general transformation function that can be applied to additional data when it becomes available (Table 1). Such a function allows researchers to “fix” a specific setting rather than have all results completely change when new data is introduced. To test the use of the learned transformation function for unseen cell types in the *source dataset* we repeated our analysis, this time holding out a complete cell type from the source set in each alignment task. We next learned the transformation based on the available data and then applied the learned function to the held out data to evaluate the batch and cell type mixing. Results are presented in Figures 5, S6, S7, S8. As the figures show, the transformation learned by SCIPR allows it to keep cell types distinct, even for the unseen source cell type, while also being able to mix the batches of unseen cell types. This is evident in the high median iLISI (1.69) and low median cLISI (1.04) scores of SCIPR-gdy on the task *PBMC:10x Chrom. (v2) A*→*10x Chrom. (v2) (CD4+ T cell held-out from source)*, where the model is fit without seeing CD4+ T cells in the source set, but is then used to transform the full source set in evaluation (Figure S8). Figure 5 displays the aligned results for the cell type not used in the learning. As the figure shows, for CD4+ and Cytotoxic T cells SCIPR-gdy is able to mix the two batches even though it had never seen these in fitting.

**Figure 5:**
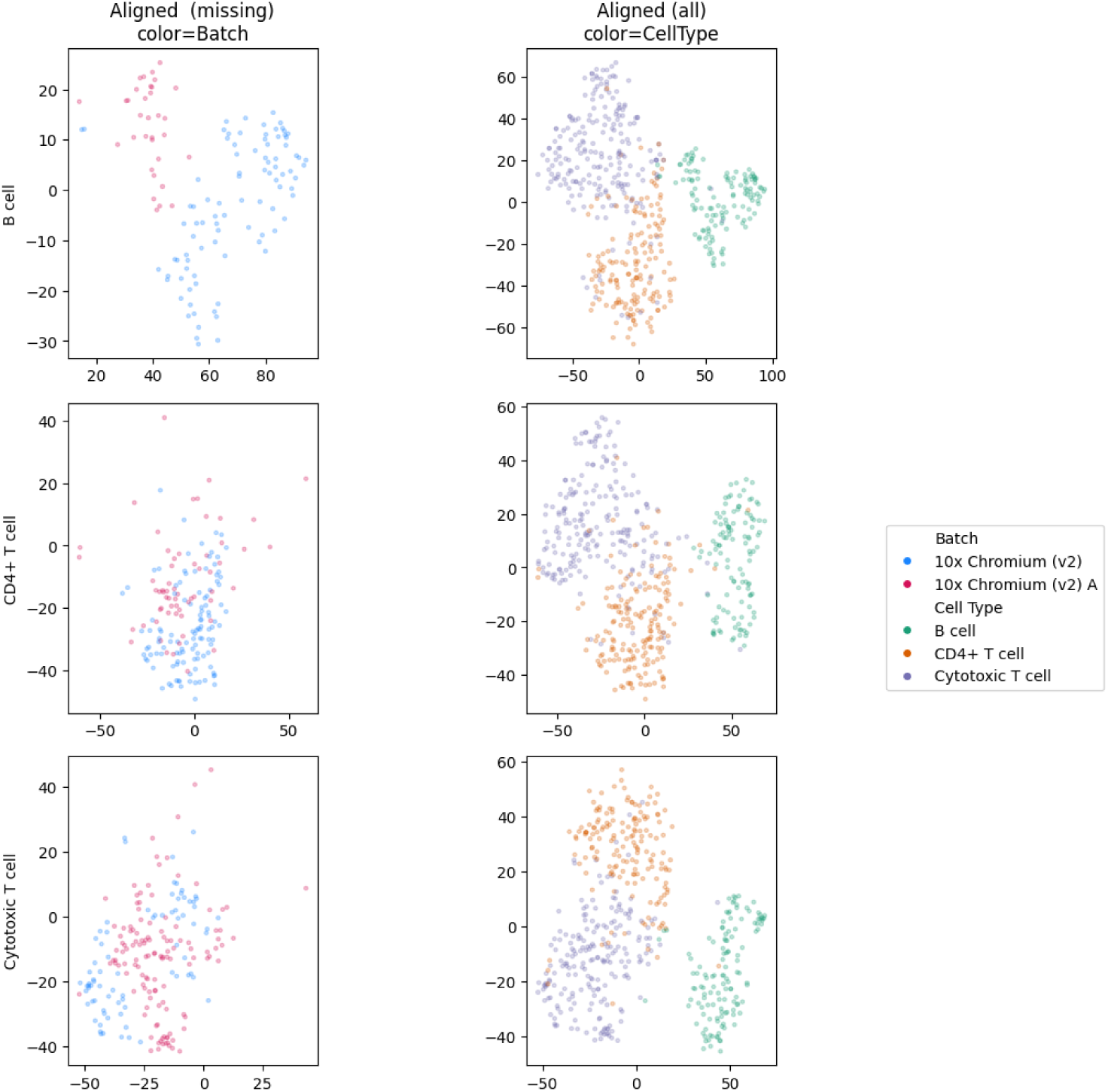
Embedding (t-SNE) visualization from the *PBMC:10x Chrom. (v2) A* →*10x Chrom. (v2)* task using SCIPR-gdy showing generalizability to new cells. In each alignment task (rows), a different cell type is completely held-out from the *source* set. The model is then fitted to align the source and the target, and the model is then used to transform the full source set, including the held-out cell type which the model did not see in the source set used for fitting. The first column shows just the held-out cell type colored by batch, after applying the fitted SCIPR-gdy model to align it. The second column shows all of the data after applying the fitted model, colored by cell type.

### 2.5 SCIPR identifies biologically relevant genes

The above results demonstrate SCIPR’s ability to integrate batches quantitatively and qualitatively. Since SCIPR achieves these results by learning a transformation function that places different weights on different genes, we next asked whether the learned weights provide information on the importance of specific genes for the set of cells being studied. To evaluate such an approach we compared ranking genes based on their SCIPR coefficients to a baseline that ranks them based on differential expression (DE) (Appendix S6, S7). Next we performed gene set enrichment analysis using the Gene Ontology to identify the significant functions associated with top genes and test their relevance (Appendix S8). The PBMC dataset, which is the largest, was also the one with the most number of significant categories identified (Table S2). When comparing top ranked genes by SCIPR and DE for the “PBMC: 10x Chrom. (v2) A → 10x Chrom. (v2)” alignment task we observed that SCIPR genes significantly overlapped with much more relevant terms when compared to DE genes for the same dataset (Table S3). For example, the top three categories for top ranked SCIPR genes are “Defense response”, “Regulation of immune response”, and “Humoral immune response” (all with adj. p-value 9.743e-9, Table S3). These categories are very relevant for blood cells given their immune system function. On the other hand, the top three categories recovered by top DE genes are much more generic and include “Ribonucleoprotein complex biogenisis”, “Ribosome assembly”, and “Cellular amide metabolic process” (Table S2).

## 3 Discussion

We presented SCIPR which extends point set registration for the alignment of scRNA-Seq data. SCIPR combines many of the desirable features of previous methods including the fact that its unsupervised, generalizable, and keeps the original (gene space) representation. Analysis of several datasets show that SCIPR successfully aligns scRNA-Seq data improving upon other methods proposed for this task. When data is missing from either the source or the target the transformation function learned by SCIPR can be used to accurately align it when it becomes available. Finally, the coefficients learned by SCIPR provide valuable information on the key genes related to the cells being analyzed.

Framing scRNA-seq alignment as a point set registration problem opens the door to applying many of the developments and advancements in that area to scRNA-seq alignment. Point set registration is a mature area that has been widely used for more than two decades. As part of this researchers looked at several different types of transformation functions, data filtration, outlier handling, and association mapping, all of which may find applications in scRNA-seq analysis.

When evaluating SCIPR and prior methods we used the local inverse Simpson’s Index (LISI) to quantify both cell type mixing and batch mixing. This leads to two values for each alignment task which can be combined for ranking the different methods by computing the difference of the medians *iLISI* − *cLISI*. Such ranking places equal weight on both issues. However, this score may not tell the whole story since some methods may be much better at one task vs. the other. For example, while SCIPR was ranked as the top method for most of the comparisons we performed, it has a tendency of to sacrifice some cell type separation in order to achieve greater batch mixing. Thus, depending on the user priorities between cell type and batch mixing, different methods may be more attractive even if the combined score is lower when compared to other methods.

While SCIPR performed best in our analysis, there are a number of ways in which it can be further improved. As mentioned above, SCIPR tends to weight batch mixing higher than cell type separation. A possible way to overcome this would be to add a regularization term to the transformation function to increase the weight of high scoring matches. Another option is to explore the use of non-linear transformations with strong customized regularization.

SCIPR is implemented as a Python package and can be downloaded from https://github.com/AmirAlavi/ scipr, and our benchmarking pipeline and data used are available at https://github.com/AmirAlavi/sc-alignment-benchmarking (see Appendix S1).

## 4 Methods

### 4.1 Dataset selection

To evaluate SCIPR and to compare its performance to previous alignment methods we used different scRNA-Seq datasets, each profiling similar cells in multiple batches. The first is the CellBench dataset (GEO: GSE118767) [25], which profiled human lung cancer cell lines and contained three batches, each from a different platform: 10x Chromium [4], Dropseq [2], and CEL-seq2 [26] (Table S1a). The smallest batch had 210 cells (Dropseq) and the largest had 895 (10x Chromium) after removing cells with low reads, and we filtered the genes to the most highly variable genes across all batches leaving us with 2351 genes (Appendix S2). The second was data from human pancreatic cells (GEO: GSE84133) [27], with four batches all using Indrop sequencing [3] where the largest batch had 1488 cells (indrop3), the smallest had 834 cells (indrop4), and we used the five largest cell types and the set of 2629 highly variable genes (Table S1b). Finally, the third and largest dataset is a PBMC dataset (GEO: GSE132044) [28] which consisted of four different batches using 10x Chromium [4] sequencing. We used the three largest cell types and the largest batch had 2510 cells (10x Chorm. (v2)), the smallest had 2011 cells (10x Chrom. (v2) A), and we used the set of 1466 highly variable genes (Table S1c). See Appendix for complete details.

### 4.2 scRNA-seq alignment

In the scRNA-seq alignment task, our goal is to learn a new representation of the data (either in the same dimensions as the original data, or in a new reduced dimension) to accomplish the following:

**Property 1** *Cell type identification: Cells from different cell types are distinct and cells from the same type are in close proximity*

**Property 2** *Batch mixing: Cells from different batches are mixed together as much as possible while respecting the first property*

### 4.3 Point set registration for single cell alignment

Unsupervised alignment of single cell data relies on the implicit assumption that the different datasets share several of the same cell types though potentially using different representations for the same type. A similar assumption is central to much of the literature in point set registration, a well-studied problem in the robotics and computer vision fields [22]. In the point set registration problem, we wish to assign correspondences between two sets of points (two “point clouds”), and learn a transformation that maps one set onto the other (Figure 1). Point sets are commonly the 2D or 3D coordinates of rigid objects, and the class of transformation function under consideration is often rigid transforms (rotations, reflections, and translations). The various point sets often originate from differing settings of sensors (viewing angle, lighting, resolution, etc) viewing the same objects or scene. Among the most widely used and classical of point-cloud registration algorithms is Bessl and McKay’s Iterative Closest Point (ICP) algorithm [23]. Briefly, each iteration of ICP has two steps: 1) assigning each point in one set (*A*, “source”) to its closest point in the other set (*B*, “target”), 2) update the rigid transformation function to transform the points in *A* as close as possible to their assigned points in set *B*. At the end of each iteration, the points in *A* are transformed via the current rigid transform and the process is repeated until convergence. Thus, each iteration of ICP can be concisely represented as minimizing the following loss function:

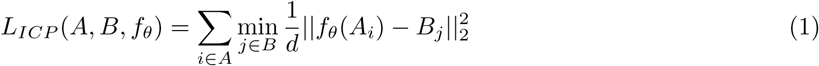

where *A, B* ∈ ℝ^*d*^, and *d* is the number of genes

and *f*_*θ*_ is further constrained to rigid transformation functions

However, applying ICP as-is to align two scRNA-seq datasets could be problematic since:

- ICP assumes that every point in *A* corresponds to a point in set *B*, whereas scRNA-seq datasets may not fully overlap in cell types. For example, in studying embyronic development, we observe the transcriptome at different embryonic days, where some cell fates exist only after a certain day [29, 30].
- ICP assumes that a rigid transform relates the two sets. This may have been appropriate for 3D rigid objects, but not for the complicated, high-dimensional single cell transcriptome data.
- ICP is prone to assigning many points in *A* to the same point in *B* (collapsing a point) even when they are not fully compatible [31]. In contrast, if the same cell type exists in both datasets we can expect the number of cells to be more balanced.

Thus, while ICP has been very successful in image analysis, it requires modifications in order to accurately align scRNA-Seq data.

### 4.4 Adapting ICP for scRNA-seq dataset alignment

Given the discussion above, both stages of ICP need to be changed in order to align scRNA-Seq data. More formally, these two stages are:

1. Assignment stage (input: point sets *A* and *B*) - assign pair set *S* ⊆ {(*i, j*) | 1 ≤ *i* ≤ |*A*|, 1 ≤ *j* ≤ |*B*|} (Figure 1 panel 2). Options for this stage may vary in the cardinality of *S*, and whether or not it allows points to be shared between pairs.
2. Transformation stage (input: assigned pairs *S*) - given *S* from the Assignment stage, learn a transform function that transforms points in *A* to reduce the mean squared error (MSE) between the assigned pairs in *S* (Figure 1 panel 3). Options for this stage vary based on the family of functions considered.

To adapt stage 1, we propose two approaches for assigning points in section 4.5: one based on a novel greedy algorithm and another based on Mutual Nearest Neighbors (MNNs). For stage 2, we set the family of transformation functions to be affine transformations (section 4.6).

### 4.5 Assigning cells between datasets

First, we focus on the Assignment stage. The input to this stage is the target set *B* and the current state of the source set of points *A* (*A* is being updated at every iteration of the algorithm). ICP computes the pairwise distance matrix between members of these sets ***D*** ∈ ℝ^*n*×*m*^ where *n* = |*A*| and *m* = |*B*| (Figure S1), and finds the element in each row with the smallest distance to match to that point in *A*.

In contrast, for scRNA-seq alignment we would like to require the following:

- Not too many points in *A* are matched with the same point in *B* (avoid collapsing many points in one dataset onto a single point).
- Not all points must be assigned (since the two dataset may not fully overlap in terms of cell types).

One approach for addressing the first requirement is using a bipartite matching algorithm [32] instead of picking the closest point. In such an algorithm a global optimal matching is found such that each point is only matched to a single point in the other set. However, such algorithms violate the second requirement since they result in “perfect” matchings, where all points in *A* are matched. An alternative is to use partial matching algorithms in which only a subset (or a fraction) of the points in A are required to be matched to points in B. Optimal partial matching is a well studied problem in the computer science literature and requires solving a min-cost flow graph problem [33]. The problem can be solved via an efficient network simplex algorithm, however, for graphs with thousands of nodes (as in single cell data) this is still rather time consuming. If we let the number of vertices be *V* = *n* + *m* (e.g. number of cells in both batches), the number of edges be *E* = *n* × *m*, and the largest edge weight (distance between points) be *C*, then the polynomial time network simplex algorithm has a run time of *O*(*V E* log *V* log(*V C*)) [34]. Given the large number of cells in each dataset such partial matching methods are too time consuming in practice (Figure S2).

Instead, in Algorithm S1 we propose an efficient greedy algorithm for partial assignment between *A* and *B*. The algorithm sorts all of the edges between members of the two sets based on distance. Next, we proceed along the ordered edges starting from the smallest distance. If an edge includes a point in set *B* that has already been selected *β* times by previously chosen edges, we discard it and continues down the list. Our algorithm has parameters to adjust how many times we allow each point in *B* to be matched to (*β*), and how many of the points in *A* must be matched (*α*). In our experiments, we set *β* = 2 and *α* = 0.5. The runtime of this algorithm is dominated by the *sortElementsAscending* function which sorts the distances leading to a worst case runtime of *O*(*E*^2^) and a much faster *O*(*E* log(*E*)) on average. Though not an optimal solution to the partial bipartite matching problem, we find that this works well in our related scRNA-seq cell pair assignment problem for alignment.

We also experiment with a matching procedure which follows the foundational work of using mutual nearest neighbors (MNNs) [16] to define our pair assignments between the two sets. For a point *i* in set *A* and a point *j* in set *B*, if *i* is in the set of *k*-nearest neighbors among *A* for point *j*, and *j* is in the set of *k*-nearest neighbors among *B* for point *i*, then *i* and *j* are MNNs. In our experiments, we set *k* = 10.

### 4.6 Learning a transformation function

So far we focused on matching points given their distance. In the next stage, we will fit a transformation function to align the matched points. As discussed above, the family of rigid transforms is not well suited to compute such alignments for scRNA-seq data (Figure S3). Instead, we propose to use the family of affine transformations to align scRNA-seq datasets. Affine transforms are of the form *f*_*θ*_(*x*) = *W*^*T*^ *x* + *b* where *θ* = {*W, b*} is the learnable weights of the function, and include rotation, reflection, scaling, and shearing.

To learn this function, we minimize an objective function that aims to move the assigned points closer to each other, via such an affine transformation. Given a pair assignment *S* from the previous step (section 4.5) (which may be the result of the classic “closest” strategy from ICP, our greedy algorithm, or MNN matching), learning the transformation function (Figure 1 panel 3) is equivalent to minimizing the loss function given as Equation 2. We note that this objective function is not over all pairs of points in sets *A* and *B*; it is computed over only those pairs of points selected in *S*, denoted by the subscript under the sum.

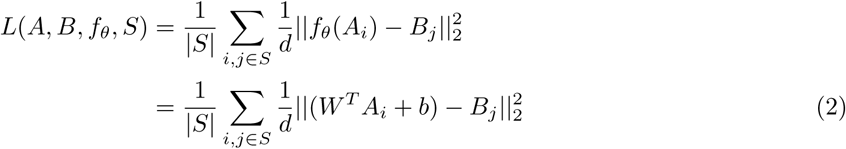

where *A, B* ∈ ℝ^*d*^, and *d* is the number of genes

This is a least-squares objective function. If the system is overdetermined, this could be solved exactly. However, due to the high dimension we are working in (each point is the expression of thousands of genes), the matrix inversion for the exact solution is expensive to compute, as matrix inversion is *O*(*d*^3^). To avoid this, we approximate the solution using gradient descent to arrive at our transformation function 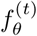 for the current iteration *t* (see Appendix S5 for gradient descent settings).

### 4.7 Iterative step

After each of the stages (assignment and transformation function update), we use our learned transformation function at the current iteration 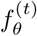 to transform the all points in *A* (not just those in the set *S* from the matching algorithm) (Figure 1 panel 4), and then repeat the stages for *T* iterations (Figure 1 panel 5). The final learned transformation of source points *A* to target points *B* is a chained series of transformations (composite function) from each iteration (Figure 1 panel 6). Since our function class for *f*_*θ*_ is affine transformations, and the composition of affine transformations is itself an affine transformation, we can combine this chain of transformations into a single affine transformation. See Appendix S4 for details.

### 4.8 Validation

A number of methods have been proposed to test the accuracy of alignment based methods [35, 24]. These evaluation metrics try to balance two, sometimes competing, attributes. The first is dataset mixing which is the goal of the alignment. The second is cell type coherence. A method that randomly mixes the two datasets would score high on the first measure and low on the second while a method that clusters each of the datasets very well but cannot match them will score high on the second and not on the first.

To track both dataset mixing and biological signal preservation, we follow [24] and use the local inverse Simpson’s Index (LISI). LISI measures the amount of diversity within a small neighborhood around each point in a dataset, with respect to a particular label. The lowest value of LISI is 1 (no diversity). As in [24], we define integration LISI (iLISI) as the score computed when using the batch label for each datapoint, and cell-type LISI (cLISI) as the score when using the cell-type label. iLISI measures the effective number of datasets within the neighborhood (so the higher the better). cLISI measures the effective number cell types within the neighborhood (so the lower the better). With these two metrics in hand, we can keep track of not only the ability of our algorithms to align one dataset onto another, but also their ability to preserve original signal. In our figures which report the iLISI and cLISI scores, we rank the methods based on the difference of medians *iLISI* − *cLISI* score to capture the ability of the emthods to maximize and minimize these two quantities respectively.

## End Notes

### Author Contributions

A.A. and Z.B.J. designed the study, the algorithms, and the analysis. A.A. developed and implemented the algorithms and performed all the analysis. A.A. and Z.B.J wrote the paper.

## Supporting information captions

- S1 Figure. The distance matrix between cells in batches *A* and *B*.
- S1 Algorithm. Greedy pair assignment algorithm.
- S2 Figure. A comparison of runtimes of a greedy matching algorithm (Algorithm S1) compared to a network flow-based approach for finding an optimal partial matching (Min Cost Flow).
- S3 Figure. Comparison of using a rigid transformation versus an affine transformation in the SCIPR method.
- S4 Figure. Quantitative scoring of alignment methods on the Pancreas dataset with a cell type held out from the target set.
- S5 Figure. Quantitative scoring of alignment methods on the PBMC dataset with a cell type held out from the target set.
- S1 Table. Cell type and batch distributions for three scRNA-seq datasets we use for evaluation.
- S6 Figure. Quantitative scoring of alignment methods on the CellBench dataset with a cell type held out from the source set.
- S7 Figure. Quantitative scoring of alignment methods on the Pancreas dataset with a cell type held out from the source set.
- S8 Figure. Quantitative scoring of alignment methods on the PBMC dataset with a cell type held out from the source set.
- S2 Table. Gene enrichment analysis of Differential Expression results.
- S3 Table. Gene enrichment analysis of model weights from SCIPR-mnn.
- S4 Table. Gene enrichment analysis of model weights from SCIPR-gdy.
- S5 Table. Quantitative alignment scores from all alignment tasks in our work.
- S1 Appendix. scRNA-seq alignment benchmarking software and data.
- S2 Appendix. Data preprocessing and filtration.
- S3 Appendix. Software and settings for related methods.
- S4 Appendix. Computing the final affine transformation at the end of SCIPR.
- S5 Appendix. Parameter settings for SCIPR experiments.
- S6 Appendix. Differential expression analysis.
- S7 Appendix. Selecting top genes from SCIPR models for enrichment analysis.
- S8 Appendix. Gene set enrichment analysis.

